# PneumoBrowse 2: An integrated visual platform for curated genome annotation and multiomics data analysis of *Streptococcus pneumoniae*

**DOI:** 10.1101/2024.08.07.606308

**Authors:** Axel B. Janssen, Paddy S. Gibson, Afonso M. Bravo, Vincent de Bakker, Jelle Slager, Jan-Willem Veening

## Abstract

*Streptococcus pneumoniae* is an opportunistic human pathogen responsible for high morbidity and mortality rates. Extensive genome sequencing revealed its large pangenome, serotype diversity, and provided insight into genome dynamics. However, functional genome analysis has lagged behind, as that requires detailed and time-consuming manual curation of genome annotations, and integration of genomic and phenotypic data. To remedy this, PneumoBrowse was presented in 2018; a user-friendly interactive online platform, which provided the detailed annotation of the *S. pneumoniae* D39V genome, alongside transcriptomic data. Since 2018, many new studies on *S. pneumoniae* genome biology and protein functioning have been performed. Here, we present PneumoBrowse 2 (https://veeninglab.com/pneumobrowse), fully rebuilt in JBrowse 2. We updated annotations for transcribed and transcriptional regulatory features in the D39V genome. We added genome-wide data tracks for high-resolution chromosome conformation capture (Hi-C) data, chromatin immunoprecipitation coupled to high-throughput sequencing (ChIP-Seq), ribosome profiling, CRISPRi-seq gene essentiality data and more. Additionally, we included 18 phylogenetically diverse *S. pneumoniae* genomes and their annotations. By providing easy access to diverse high-quality genome annotations, and links to other databases (including UniProt and AlphaFold), PneumoBrowse 2 will further accelerate research and development into preventive and treatment strategies, through increased understanding of the pneumococcal genome.

**Graphical abstract:** 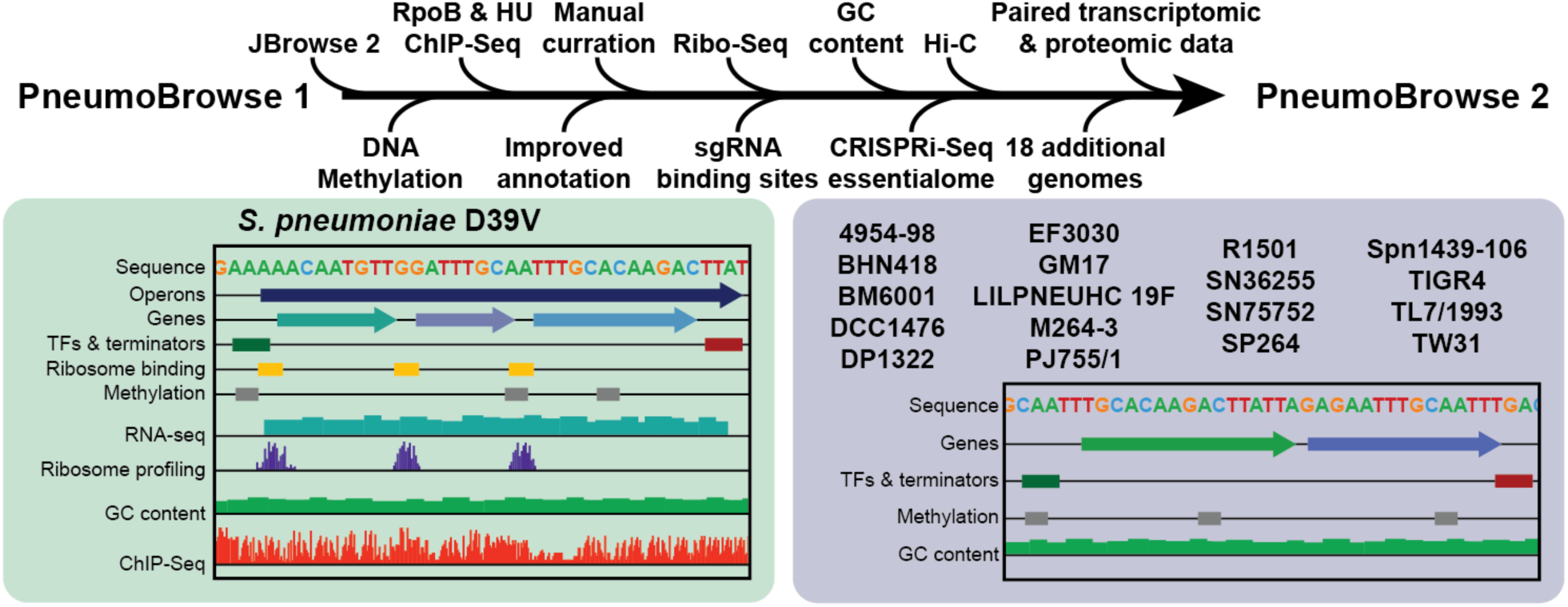

## Introduction

*Streptococcus pneumoniae* is a clinically highly relevant opportunistic human pathogen responsible for over 100 million episodes of lower respiratory tract infections that result in over 500.000 deaths annually. Simultaneously*, S. pneumoniae* is a harmless constituent of the nasopharyngeal microbiome in a large part of the human population (1–3). The balance between lethal opportunistic pathogen or harmless nasopharyngeal commensal is a product of the genomic content and its expression. How the delicate balance between these two lifestyles is controlled is an active area of research. Improved comprehension may come from a more detailed understanding of the regulatory mechanisms surrounding the expression of its genomic contents. Visualizing the genome and its contents, along with data from regulatory and expression studies, can reveal new insights that might otherwise have gone unnoticed.

To understand the gene expression and regulatory networks of *S. pneumoniae*, the oft-used serotype 2 strain D39 (4) and its derivatives such as R6, D39V and D39W have been subjected to extensive DNA and transcriptome profiling studies (5–11). This has resulted in a detailed understanding of the genomic sequence, operon structure, regulatory networks, and transcriptome in response to a variety of conditions. These results were bundled in PneumoBrowse (referred to as “PneumoBrowse 1” from here), which was launched in 2018 (5). PneumoBrowse 1 provided an intuitive, user-friendly, and visual platform through which the highly detailed manual annotation, including coding features, predicted terminator sequences, and repeat sequences, of the circular genome sequence of *S. pneumoniae* D39V could be inspected, but also the in-depth transcriptomic sequencing data. PneumoBrowse 1 thus provided a valuable platform for the pneumococcal research community to gain an in-depth understanding of the genome contents and expression of this important model strain.

Since the release of PneumoBrowse 1, numerous subsequent studies further delved into the genome biology of *S. pneumoniae* using state-of-the-art methods, thereby revealing new complexities in its genome biology. One recently developed method to interrogate pneumococcal gene function on a genome-wide basis has been CRISPR interference (CRISPRi) coupled to next generation sequencing (CRISPRi-Seq) (12, 13). CRISPRi libraries have been used to determine genome-wide essentiality under different conditions ((12, 13) and de Bakker and Veening, under review). Additional genome-wide works have studied translated features through ribosome profiling (also known as ribosome footprinting, or Ribo-Seq) (14); the correlation between RNA and protein abundances through transcriptomics and proteomics (de Bakker and Veening, under review); as well as chromosome conformation and protein binding through high-resolution chromosome conformation capture (Hi-C) and chromatin immunoprecipitation coupled to high-throughput sequencing (ChIP-Seq) experiments (Mazzuoli, van Raaphorst, and Veening, under review).

Here, we introduce PneumoBrowse 2, to include the newly available data, but also several major updates. First, we updated the underlying software to JBrowse 2 (15), which vastly improves user experience whilst simultaneously increasing the platform capabilities. Second, we have included the results of the genome-wide Hi-C, ChIP-seq, transcriptomic, proteomic, and ribosome profiling studies in D39V. We have also added the binding site locations of the sgRNAs in the CRISPRi library used in recently published work, as well as the gene essentialities determined under different conditions using this library (12, 13). In addition, we manually curated and updated gene annotations based on recent molecular studies, and added direct links for each transcribed feature to facilitate the use of other specialized databases, such as UniProt and AlphaFold.

Besides updates to the D39V annotation, we also introduce the genomes of 18 other, phylogenetically diverse, pneumococcal strains in PneumoBrowse 2. Although *S. pneumoniae* strain D39 (derived strains) are among the most investigated strains, other strains are also used to explore pneumococcal biology due to their diverse traits. For example, TIGR4 was the first pneumococcal strain to be entirely sequenced (16), whilst strains BM6001 and DP1322 are used to investigate integrative conjugative element (ICE) biology because they ICEs encoding several antibiotic resistance genes (17, 18). Strain BHN418 is known for its use in human challenge models (19, 20), whilst strains EF3030 and LILPNEUHC 19F are used to investigate the clinically relevant serotype 19F group of *S. pneumoniae* (21). We include the long-read assembled genomes of these and other, phylogenetically diverse, *S. pneumoniae* strains in PneumoBrowse 2. These genomes have been annotated for genomic content, transcription factor bindings sites, Rho-independent terminator sequences, and chromosomal DNA methylation patterns.

With these improvements, PneumoBrowse 2 will serve as a portal to the latest insights on the contents and workings of a wide variety of pneumococcal genomes. By making these data accessible for a wide audience, we anticipate that the pneumococcal research field can make additional progress into the biology of these diverse *S. pneumoniae* strains. PneumoBrowse 2 is freely available on https://veeninglab.com/pneumobrowse.

## Methods

### Strains used in this work

The strains included in PneumoBrowse 2 are listed in Table 1.

**Table 1.** S. pneumoniae strains presented in PneumoBrowse 2. . The strain names used in PneumoBrowse 2, alternative identifiers(s), multilocus sequence type, serotype, and accession codes for the genomic and sequencing data deposited at the Sequence Read Archive (SRA). The origin of the strain is noted if no previous reference is available. Strains that serve as a reference clone for one of the 43 clones defined by the Pneumococcal Molecular Epidemiology Network (https://www.pneumogen.net/gps/#/resources#pmen clones) are indicated as such. No MLST type could be defined for strains BM6001 and DP1322 through mlst. Strains DP1322 and R1501 are derivatives of S. pneumoniae strain R6, and do not encode a functional capsular polysaccharide biosynthesis gene cluster in their genomes, and are thus acapsular. The analysis of PacBio sequencing reads for methylated bases through SMRT Link did not yield results for strains R1501 (only bases with unspecified tag “modified_base”) and SN75752 (none of the SMRT Link called hits passed quality filtering parameters). NA; not applicable.

| Pneumobrowse ID | Alternative IDs | MLST | Serotype | SRA Accession | Genome Accession | Origin | Reference Strain | Remark | Reference |
| --- | --- | --- | --- | --- | --- | --- | --- | --- | --- |
| 4954-98 | Tennessee14-18, ATCC BAA-340 | 67 | 14 | SRR28781549 | JBDIMH000000000 | NA | Yes; PMEN18 reference strain |  | Gherardi et al., Journal of Infectious Diseases, 2000 |
| BHN418 | NA | 138 | 6B | SRR28790986 | CP155538 | NA | No |  | Browall et al., Journal of Infectious Diseases, 2014 |
| BM6001 | NA | Unknown | 19F | SRR28781558 | JBDIMJ000000000 | NA | No | Carries Tn5253 composite integrative conjugative element. | Vijayakumar et al., Journal of Bacteriology, 1986 |
| D39V | NCTC 14078 | 595 | 2 | See Slager et al., Nucleic Acids Research, 2018 | See Slager et al., Nucleic Acids Research, 2018 | NA | No | NCTC 7466 derivative | Slager et al., Nucleic Acids Research, 2018 |
| DCC1476 | Sweden15A-25, ATCC BAA-661 | 156 | 15A | SRR28781547 | CP155534 | NA | Yes; PMEN25 reference strain |  | Sá-Leão et al., Journal of Infectious Diseases, 2000 |
| DP1322 | NA | Unknown | Acapsular | SRR28781557 | CP155537 | NA | No | R6 (acapsular D39 derivative) derivative carrying Tn5253 | Vijayakumar et al., Journal of Bacteriology, 1986 |
| EF3030 | NA | 43 | 19F | SRR28781559 | JBDIMI000000000 | NA | No |  | Andersson et al., Journal of Experimental Medicine, 1983 |
| GM17 | 700670 | 90 | 6B | SRR28781552 | CP157351 | NA | Yes; PMEN2 reference strain |  | McGee et al., Journal of Clinical Microbiology, 2001 |
| LILPNEUHC 19F | NA | 179 | 19F | SRR28781556 | JBDIMK000000000 | Blood isolate from patient with invasive pneumococcal disease. | No |  | This study |
| M264-3 | Netherlands3-31, ATCC BAA-1657 | 180 | 3 | SRR28781545 | CP155536 | NA | Yes; PMEN31 reference strain |  | Enrich and Spratt, Microbiology, 1998 |
| PJ755/1 | Sweden1-28, ATCC BAA-1655 | 306 | 1 | SRR28781546 | CP155535 | NA | Yes; PMEN28 reference strain |  | Henriques-Normark et al., Journal of Infectious Diseases, 2001 |
| R1501 | NA | 5954 | Acapsular | SRR29015166 | CP155800 | NA | No | R6 derivative. No modified bases resulted from analysis of the PacBio sequencing data due to insufficient quality of SMRT Link-hits. | Dagkessamanskaia et al., Molecular Microbiology, 2004 |
| SN36255 | NA | 320 | 19A | SRR28781561 | CP155531 | Blood isolate from patient with invasive pneumococcal pneumonia. | No |  | This study |
| SN75752 | NA | 6521 | 11A | SRR28781553 | see Gibson et al., Plos Pathogens, 2022 | NA | No | Referred to as "11A" in reference. No modified bases resulted from analysis of the PacBio sequencing data due to insufficient quality of SMRT Link-hits. | Gibson et al., Plos Pathogens, 2022 |
| SP264 | Spain 23F-1, ATCC 700669 | 81 | 23F | SRR28781560 | CP155532 | NA | Yes; PMEN1 reference strain |  | McGee et al., Journal of Clinical Microbiology, 2001 |
| Spn1439-106 | Colombia 5-19, ATCC BAA-341 | 289 | 5 | SRR28781548 | CP155533 | NA | Yes; PMEN19 reference strain |  | Tamayo et al., Journal of Clinical Microbiology, 1999 |
| TIGR4 | NA | 205 | 4 | SRR28781554 | CP155539 | NA | No |  | Gibson et al., Plos Pathogens, 2022 |
| TL7/1993 | Spain 9V-3, ATCC 700671 | 156 | 9V | SRR28781551 | CP156625 | NA | Yes; PMEN3 reference strain |  | McGee et al., Journal of Clinical Microbiology, 2001 |
| TW31 | Taiwan19F-14, ATCC 700905 | 236 | 19F | SRR28781550 | JBDIMG000000000 | NA | Yes; PMEN14 reference strain |  | McGee et al., Journal of Clinical Microbiology, 2001 |

### PneumoBrowse 2 web interface

PneumoBrowse 2 is available online (https://veeninglab.com/pneumobrowse) through JBrowse 2 (15). The NucContent plugin for JBrowse2 is used to provide a track to calculate and visualize GC content (available from https://github.com/jjrozewicki/jbrowse2-plugin-nuccontent).

### Isolation of genomic DNA

Genomic DNA was isolated through previously published methods, with minor adaptations (22, 23). Bacterial cultures were grown to an optical density at 595 nm of 0.2 in C+Y medium without shaking, spun down, and resuspended in Nuclei Lysis solution containing 0.05% SDS, 0.025% deoxycholate, and 200 μg/mL RNase A. The suspension was incubated at 37°C (20 minutes), 80°C (5 minutes), 37°C (10 minutes), and cooled to room temperature. Lysates were treated with Protein Precipitation Solution, vortexed vigorously, incubated on ice for 10 minutes, then pelleted by centrifugation. DNA was precipitated from supernatants with isopropanol and collected by centrifugation. Pellets were washed once in 70% ethanol, air-dried, and resuspended in water.

If isopropanol failed to produce sufficient DNA, FastPure DNA Isolation Mini Kit columns (Vazyme) were used. After protein precipitation and centrifugation, supernatant was transferred to the column and processed according to protocol. DNA was eluted off the column in water.

### Whole genome sequencing and assembly

DNA concentrations were quantified with a Qubit 4.0 fluorometer, using the high-sensitivity double-stranded DNA kit. Quality control for chromosome integrity was performed by running the gDNA on a 1% agarose gel, and through an Agilent Fragment Analyzer.

Long-read sequencing for 4954-98, BM6001, DCC1476, DP1322, EF3030, GM17, LILPNEUHC 19F, M264-3, PJ755/1, R1501, SN36255, SP264, Spn1439-106, TL7/1993, and TW31 was performed using a PacBio Sequel II machine. PacBio reads were demultiplexed, filtered for quality, and assembled through SMRT Link (24).

Long-read sequencing for BHN418 was performed using an Oxford Nanopore MinION Mk1B, the Rapid Sequencing Kit V14, and an R10.4.1 flow cell. Nanopore POD5 files were base-called using Dorado (v0.6.0; available from https://github.com/nanoporetech/dorado) using the “super accurate basecalling model”. Resulting FASTQ files were used for assembly using Unicycler (v0.5.0) (25).

### Genome annotation and characterization

Assembled genomes were annotated through Prokka (v1.14.6) (26). The Prokka script was adapted by removing the “-c” flag from Prodigal (27), to enable annotation of open reading frames without a start or stop codon at the ends of a contig. Aragorn (v1.2.38) was used for the annotation of transfer RNAs (28); Infernal (v1.1.4) and Rfam were used to annotate non-coding RNAs (29); Minced (v0.4.2) was used to identify CRISPR arrays (30); and HMMER3 (v3.3.2) was used for protein similarity searching (31). The previously produced high-detail annotation of D39V was used as reference (5). Rho-independent terminators were predicted through TransTermHP (v2.09) (32), and filtered for a minimal score of 60. ComE, ComX, and ParB binding sites were determined through FIMO (MEME suite v5.5.5), using the previously determined consensus sequences in *S. pneumoniae* D39V (5, 7, 33, 34), and filtered based on their q-values (q =< 0.001 for ComE and ComX; q =< 0.01 for ParB).

Modified bases were called from the PacBio reads using SMRT Link and filtered for a QV score >= 100. (Hydroxy)methyl modifications were determined from Nanopore POD5 reads using Dorado (v0.6.0) using the “super accurate model”, extracted using modkit (v0.2.7; available from https://github.com/nanoporetech/modkit), and filtered based on the number (>=20), and fraction (>=95%) of reads supporting the modification.

Additional functional annotation for the D39V genome was performed using the online eggNOG-mapper (v2) tool (35–39).

MLST typing for all genomes was performed using mlst (v2.23.0; available from https://github.com/tseemann/mlst).

### Genome comparison, and core genome phylogenetic analysis

Closely related genomes were aligned using the command-line version of progressiveMauve (version of February 13, 2015) (40).

A core genome alignment for all genome sequences included in PneumoBrowse 2 and genomes representing the phylogeny of *S. pneumoniae* was built using snippy (v4.6.0, available from https://github.com/tseemann/snippy), using D39V as reference (41). Duplicate genomes were filtered out of the genome set from Antic *et al..* Alignments were corrected for recombination using Gubbins (v3.3.1) (42). The phylogeny was constructed using FastTree (v2.1.10) (43), and mid-point rooted and visualized using FigTree (v1.4.5) (available from http://tree.bio.ed.ac.uk/software/figtree/).

### Data availability

All long-read sequencing data and base modification files for PacBio sequenced genomes (except for D39V), are available in the NCBI Sequence Read Archive under BioProject accession code PRJNA1103744. Assembled genomic sequences are available from NCBI through the respective accession codes (Table 1). Genome sequence and methylation data from D39V are available under BioProject accession code PRJNA295913 (5). The SN75752 genomic sequence is available under accession code CP089949.1 (44).

## Results

### Upgrading PneumoBrowse to JBrowse 2

PneumoBrowse 1 was set up using the JBrowse 1 software (45). However, the halted development of JBrowse 1 makes PneumoBrowse 1 increasingly harder to maintain and to update with new data due to outdated libraries and plugins. To remedy this, we updated the underlying software running PneumoBrowse to the newly developed JBrowse 2, which offers significant upgrades compared to JBrowse 1 (15). The upgrade ensures the longevity of PneumoBrowse 2 by accommodating a broader range of data types and offering simplified methods for adding them, making future maintenance more efficient.

The update to JBrowse 2 also provides a significant increase in quality of experience for users. For example, data tracks are now loaded faster, and users may share, restore, or export their session, which will preserve the user’s current view for future reference. Thus, PneumoBrowse has had a significant increase in performance, user experience, and flexibility for future updates with the update to JBrowse 2.

### Further detailing the annotation of the *S. pneumoniae* D39V genome

Since the introduction of PneumoBrowse 1, new studies have further explored the genome biology of *S. pneumoniae*. To reflect the state-of-the-art in pneumococcal genome biology, we have further curated the existing detailed manual annotation of the *S. pneumoniae* D39V. In total, 623 changes to features in the annotation of D39V were made.

Out of these 623 changes, 375 changes were made to the annotations of transcribed features (Supplemental Table S1). The genomic annotation now reflects the latest insights in the genome of *S. pneumoniae* through 230 changes to previously existing annotations of transcribed features. For example, the annotation for SPV_0476 has been updated to code for the pneumococcal spatiotemporal cell cycle regulator CcrZ, and SPV_0878 has been revised to encode the regulator of chromosome segregation RocS respectively, since the products of these genes have now been shown to play crucial roles in the pneumococcal cell cycle (22, 46). Another study has annotated a group of aquaporins at loci SPV_1320 (*Pn-*aqpC), SPV_1569 (*Pn-aqpA*), and SPV_2011 (*Pn-aqpB*) (47). In addition, the functions of SPV_1416 (*murT)* and SPV_1417 (*gatD*) in peptidoglycan biosynthesis have been determined (13).

Besides modifying previously existing annotations, we have also added 122 new locus tags to reflect the annotation of previously unknown transcribed features in the D39V genome. Of these 122 locus tags, 114 (SPV_2459 - SPV_2572) are small open reading frames (sORFs), that are the result of the identification of their previously unannotated translational start sites through ribosome profiling (14). These sORFs are now annotated as Ribo-Seq-identified ORF genes (*rio* genes) (Figure 1A, Supplemental Table S2). Four *rio* genes were further annotated: *rio9* and *rio12* were annotated as *rtgS1* and *rtgS2* (48), *rio82* as *shp1518* (49), and *rio3* was shown to be important for nasopharyngeal colonization, although the biological mechanism remains unclear (14). The remaining 110 *rio* genes currently remain without a known biological function. Four additional transcribed features in the *rtg* locus containing peptidase ABC-transporters (SPV_2573 - SPV_2576) (48), and four non-coding RNAs with unknown features propagated from the TIGR4 annotation in RegPrecise (SPV_2577 - SPV_2580) (50) have also been newly annotated. In addition, the exact coordinates of 17 transcribed features were updated, 15 of which were updated through inspection of the ribosome profiling data and cross-referencing with other databases (14, 51). Finally, *srf-01* was changed in coding feature type (from ncRNA to pseudogene) (7), and ncRNA *srf-06* (SPV_2120) was replaced by *shp144* (SPV_2573) (52, 53).

**Figure 1.**
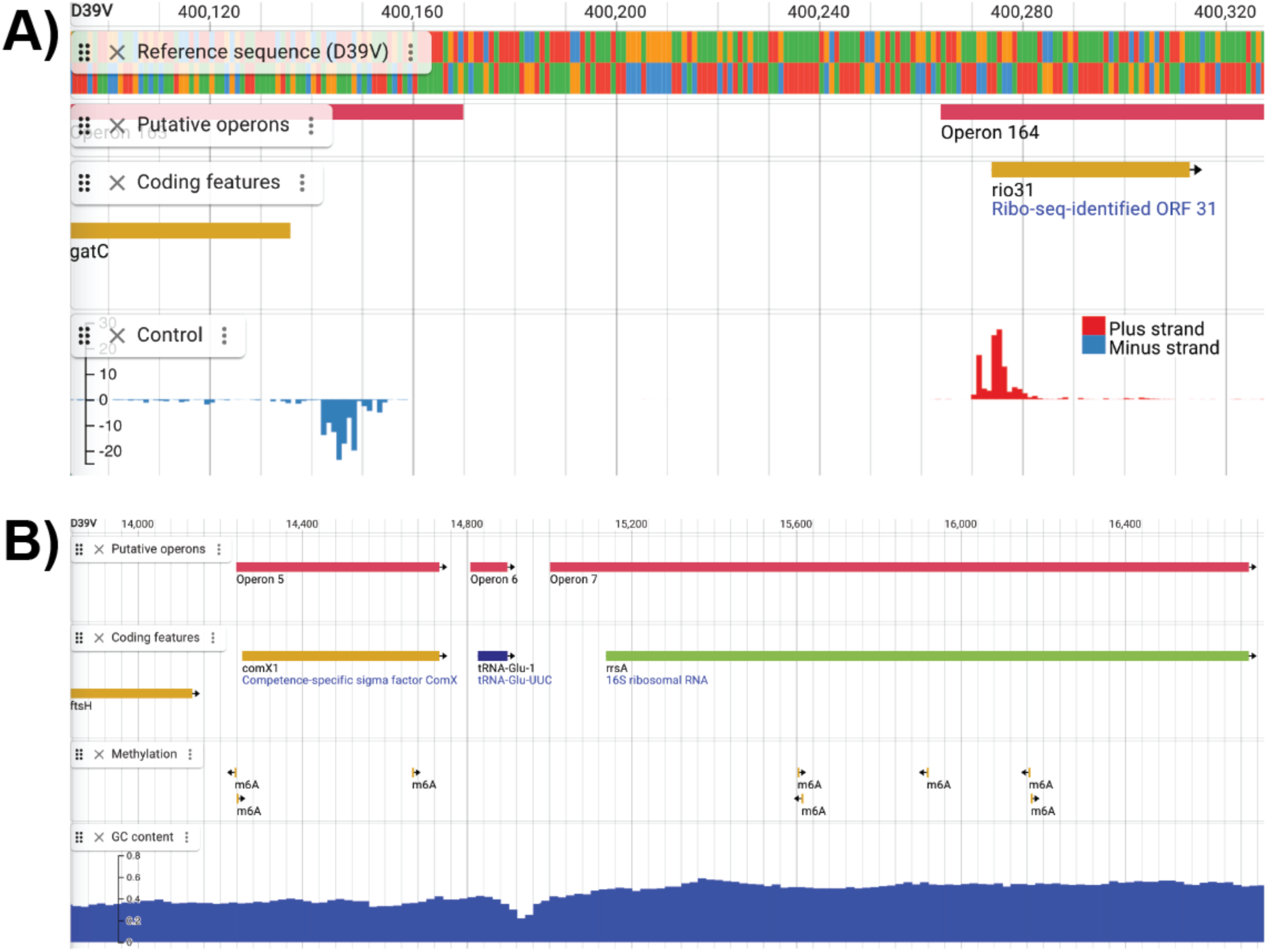
Genome-wide data tracks for D39V in PneumoBrowse 2. **A)** A genome-wide data track for the normalized intensities of ribosome profiling under control conditions has been added for D39V (14). The data track includes the score for the positive strand (upwards, red), and the negative strand (downwards, blue). **B)** Genome-wide data tracks for methylated bases in the chromosome of D39V, and GC content. The annotations of the methylated sites indicate the type and strand-specificity of the methylated bases. A track for the calculation of GC content (using customizable settings) is provided through the NucContent plugin.

Besides 374 changes to transcribed features, 249 changes are made to annotations of features involved in transcriptional regulation, such as transcription start sites, −10 signals, −35 signals, and transcription factor binding sites (Supplemental Table S2). A total of 60 annotations were added for transcription factor binding sites, including for those transcription factors crucially involved in competence and transformation, such as ComE and ComX (6, 54). Moreover, 31 previously existing transcription factor binding site annotations were modified for their exact location and binding moiety (7, 54). Further refinement of transcriptional regulation annotations came from the addition of 104 −10 signals, −35 signals and transcriptional start sites to the D39V annotation through either manual curation of RNA-seq data, or from transcriptional landscape studies of *S. pneumoniae* (7, 54). In contrast, 54 annotations previously made through algorithmic analysis have been removed after manual visual inspection of RNA sequencing data (e.g., no appreciable signal in 5’-enriched RNA sequencing coverage or located within repeat regions).

### Addition of genome-wide data tracks for D39V

Individual annotations in the D39V genome give detailed annotation about the encoded contents of the genomic sequence. However, this does not give insight into the overall regulation and expression of the genome. Besides the binding of transcription factors, gene expression is also influenced through the methylation of chromosomal DNA (55). In PneumoBrowse 2, we have now added a data track to show the position of the methylated bases in the D39V genome, determined previously from PacBio sequencing reads (Figure 1B) (5, 55). This resulted in the annotation of 3655 N^6^-methyladenosine bases and three N^4^-methylcytosine bases. For each modified base, the QV score of the call is also available.

The binding the RNA polymerase, which consists of five subunits, including the β subunit RpoB, can be influenced by binding of different transcriptional regulators to the chromosome. The binding of RpoB thus indicates sites of active transcription. In addition to RpoB binding, HU (the pneumococcal histone-like protein, also known as HlpA) can influence transcription and replication through its role in chromosomal organization by facilitating condensation. The sites to which RpoB and HU bind to the chromosome of exponentially growing *S. pneumoniae* D39V at 37°C in C+Y medium, have now been determined using ChIP-Seq studies (Mazzuoli, van Raaphorst, and Veening, under review). The data from these ChiP-Seq studies are presented in PneumoBrowse 2 as the fold enrichment of immunoprecipitated samples compared to controls (Figure 2A).

**Figure 2.**
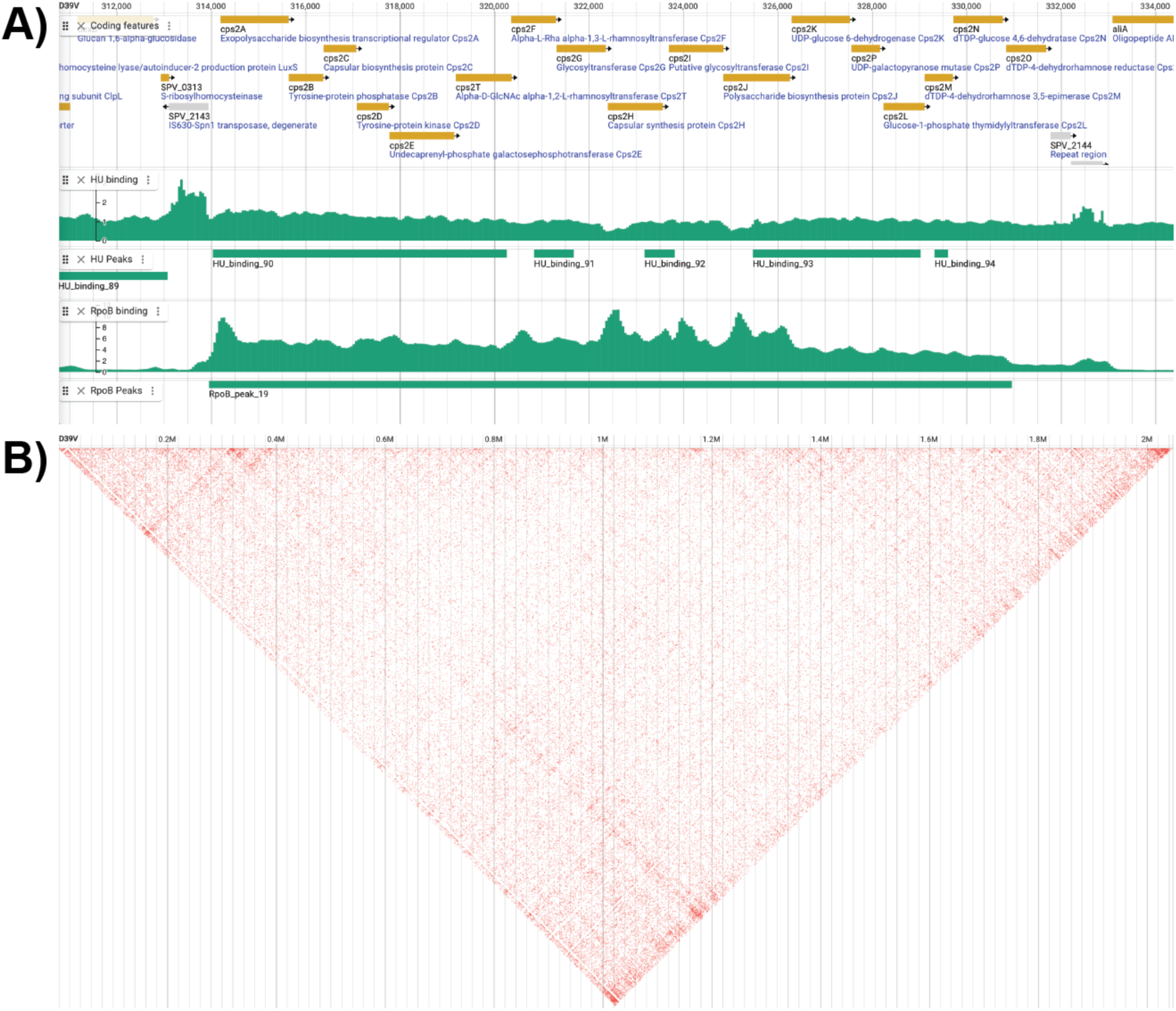
Tracks with ChIP-Seq data for chromosomal protein binding, and Hi-C conformation capture data in PneumoBrowse 2. **A)** Genome-wide data tracks representing ChIP-Seq results for HU and RpoB binding to the chromosome. The plotted values indicate the ratio of the signal between the immunoprecipitated and the control experiments, at a resolution of 50 bp. MACS2-called peaks in the RpoB- and HU-binding data indicate regions of enriched binding (Mazzuoli, van Raaphorst, Veening, under review). **B)** Hi-C chromosome conformation capture data with a resolution of 1000 bp shows the interaction of different parts of the chromosome on a diagonal line, with interactions marked in red (Mazzuoli, van Raaphorst, Veening, under review).

Although elucidation of the binding locations of HU through ChIP-Seq studies may indicate the local condensation state of the chromosome, Hi-C experiments are needed to determine which parts of the genome are in close proximity to each other, to define the overall chromosome architecture (56). The novel support of JBrowse 2 data from these types of experiments now allows us to present data on the chromosomal architecture of *S. pneumoniae* D39V (Figure 2B) (Mazzuoli, van Raaphorst, and Veening, under review). PneumoBrowse 2 presents the Hi-C data of D39V through a contact heat map in which regions that are observed to be in contact are shaded on the diagonal.

Finally, we have also added a separate track to display the GC content of the genome (Figure 1B). Visual inspection of the GC content can highlight recently horizontally acquired regions such as integrative conjugative elements or prophages (57). The calculation of the GC content can be adapted through the settings for the NucContent plugin by setting the window size (in bp), window overlap (in percentage), and the calculation method (average, or skew).

### Gene essentiality and expression data under infection(-mimicking) conditions

Important outstanding questions in pneumococcal biology include not only how the genome is expressed during infection conditions, but also which genes are essential under such conditions. As the nutrients available to the bacterial cells will differ between growth conditions (e.g., niches in the human body, or growth media), gene essentiality will also differ between conditions. To clarify which genes are essential under which conditions, a genome-wide single guide RNA (sgRNA) library in which every known operon is targeted by an sgRNA, was developed for the D39V genome. For each of the 1498 unique sgRNAs this library, the binding site position is now available in a separate genome-wide data track (Figure 3A) (12, 13).

**Figure 3.**
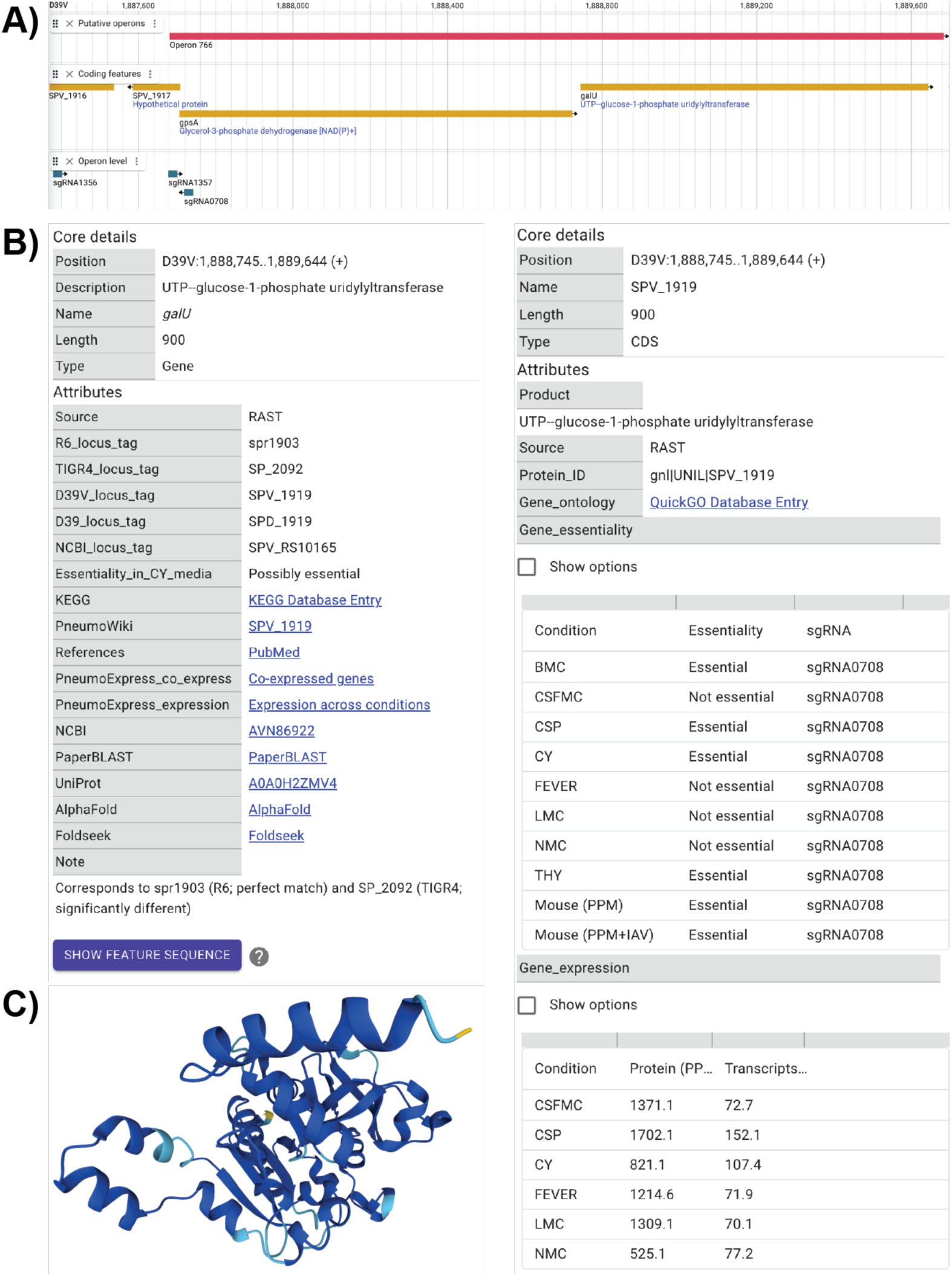
Position of sgRNAs binding sites and detailed information of annotated features for the D39V genome in PneumoBrowse 2. **A)** A genome-wide data track displaying the position of the sgRNAs included in the CRISPRi-Seq library for D39V (12, 13). The positions of the 20-nucleotide long sgRNA spacers are on the non-template strand of the targeted coding feature. **B)** Overview of the information available for each coding feature in the D39V annotation: the information for *galU* (SPV_1919) is shown. Information for each coding sequence includes links to external resources (AlphaFold, PneumoWiki, UniProt, amongst others), gene essentiality data, and expression data (12) (de Bakker and Veening, under review). **C)** The predicted AlphaFold structure of GalU that is available through the AlphaFold and UniProt links in PneumoBrowse, is shown.

Using CRISPRi-Seq, this sgRNA library has been used to determine gene essentiality in a murine pneumococcal pneumonia model (PPM), a murine pneumococcal pneumonia model with an influenza A virus superinfection (PPM + IAV), and under eight *in vitro* (infection-mimicking) growth conditions ((12), de Bakker and Veening, under review). For each transcribed feature, the essentiality under these conditions is given as either “essential” or “non-essential”, per sgRNA targeting that feature (Figure 3B).

In addition, gene essentiality under *in vitro* infection-mimicking growth conditions, coupled transcriptomic and proteomic experiments have determined the transcriptional and translational activity for each coding feature under six distinct *in vitro* (infection-mimicking) growth conditions (de Bakker and Veening, under review). In PneumoBrowse 2, these data are added as transcripts per million (TPM) or proteins per million (PPM) to reflect their relative abundance (Figure 3B).

As previously noted, the use of ribosome profiling has led to the annotation of 122 new coding features in D39V. To clarify the annotation origin of these locus tags, we have included the ribosome profiling data that demonstrates the sites of active translation in three new genome-wide data tracks (14). These data contain tracks for control experiments, but also experiments in which retapamulin and lefamulin were used. Although normal conditions may already elucidate the position of the ribosome upon initiation of translation, the addition of retapamulin and lefamulin, two pleuromutilin-class antibiotics that stall translation and thus halt the ribosome after it has bound to an mRNA molecule, further aids the identification of previously unidentified start codons. The data tracks display the genome-wide normalized abundances of the RNA sequences to which ribosomes were bound. These values are displayed as mapping to the positive (red peaks; upwards) or negative (blue peaks; downwards) strands of the D39V genome (Figure 1A), according to the identified sequences.

### Improving gene function annotation by cross-referencing with PneumoWiki, UniProt, and AlphaFold

The addition of genome-wide data tracks provides useful additional information about the expression and regulation of annotated features. However, these data cannot provide functional information of those features. To provide the best functional annotation possible, we have enriched the information for each coding feature with data from recent publications and several external databases. For example, the information for each gene now includes data on its expression and regulation (when available; Figure 3B), but also the corresponding locus tags in other *S. pneumoniae* strains (D39 (SPD_XXXX), R6 (sprXXXX), and TIGR4 (SP_XXXX)) and the NCBI locus tag (RS_SPVXXXXX). In addition, we also provide cross-referencing to other databases: including KEGG and QuickGo (both extracted from eggNOG 5.0), PneumoWiki (https://pneumowiki.med.uni-greifswald.de), PneumoExpress (both expression, and co-expression data; (6)), Foldseek (58), the NCBI, UniProt, (51), and AlphaFold (59, 60) (Figure 3C). For translated features, we provide a direct link to PaperBLAST using amino acid sequences (61).

### Expanding PneumoBrowse 2 with 18 additional genomes from phylogenetically diverse strains

We have taken advantage of the multi-genome capability of JBrowse 2 by adding the genomic sequences of 18 other *S. pneumoniae* strains to PneumoBrowse 2 (Table 1). We performed long-read sequencing for the genomes of several oft-used strains, e.g. EF3030 (serotype 19F), TIGR4 (serotype 4), and BHN418 (serotype 6B), but also for reference strains of several clones of the Pneumococcal Molecular Epidemiology Network (PMEN) (https://www.pneumogen.net/), including penicillin non-susceptible reference strains SP264 (PMEN18, serotype 14) and TW31 (PMEN14, serotype 19F). These genomes represent a phylogenetically diverse selection of *S. pneumoniae* strains (Figure 4A). For each of these 18 additional genomes, we provide the long-read assembled genomes in PneumoBrowse 2, as well as their genomic annotations, predicted transcriptional regulators binding sites, Rho-independent terminator sequences, and genomic base-modifications (Figure 4B). Within the annotation, enzyme classifiers (e.g. Prodigal classifiers and Enzyme Commission numbers), and links to AlphaFold, Foldseek, and Uniprot are provided.

**Figure 4.**
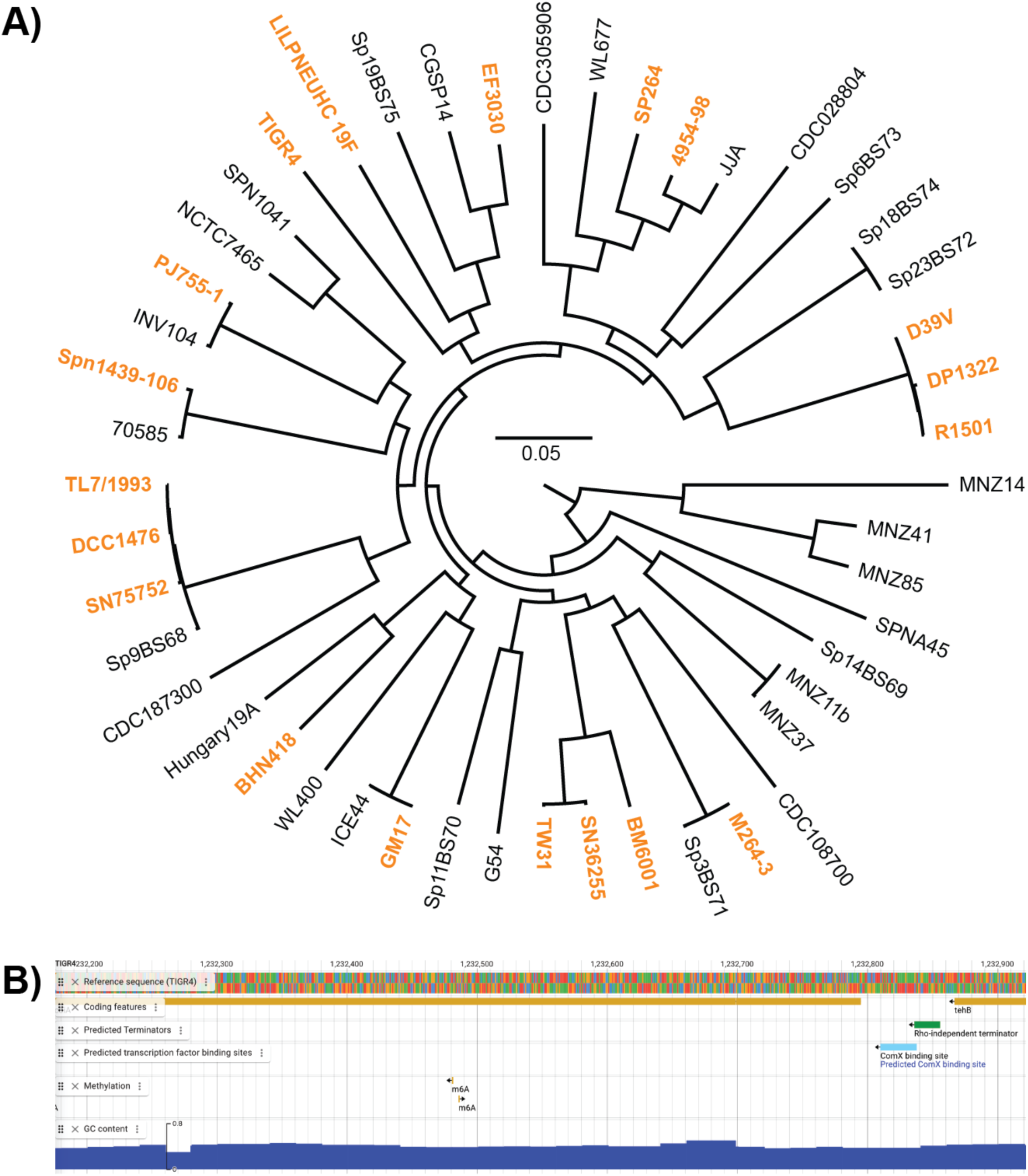
Addition of 18 phylogenetically diverse pneumococcal genomes besides D39V to PneumoBrowse 2. **A)** A midpoint-rooted maximum likelihood phylogenetic tree based on the core genome alignment of the genome sequences presented here, and genomes representing the phylogenetic diversity within the *S. pneumoniae* species, based on a recombination-corrected core genome alignment (41). Genomes included in PneumoBrowse 2 are highlighted in orange boldface text. **B)** For the 18 genomes besides D39V, tracks describing the following are available: the reference sequence; genomic annotation; predicted Rho-independent terminator sequences; predicted binding sites for ComE, ComX, and ParB; and methylated genomic bases (not available for SN7572 and R1501). A track for the calculation (using customizable settings) of GC content is provided through the NucContent plugin.

Although the strains presented in PneumoBrowse 2 are of a diverse phylogenetic standing within the *S. pneumoniae* species, some strains are observed to cluster within the same branch (Figure 4A). For example, the close clustering of D39V, DP1322, and R1501 is noteworthy, but is explained by their pedigree, as they originate from the D39 strain (4), which is a common model-strain for pneumococcal pathogenesis. The loss of the capsule locus by D39 led to the definition of the R6 branch, which later led to the creation of strain DP1322 (encoding Tn*5253*) and strain R1501 (harboring a *comC* deletion) (62). Prolonged cultivation and spread of D39 led to the definition of plasmid-free D39V (5). Likewise, the genomes of TL7/1993, DCC1476, and SN75752 (serotypes 9V, 15A and 11A, respectively) on one hand, and TW31 and SN36255 (serotypes 19F and 19A) on the other hand are also observed to cluster together in two distinct branches. As strains SN3625 and SN75752 were isolated from infections of hospitalized patients, their likeness to reference strains for clinically relevant PMEN clones is not surprising. The other genomes are observed to be phylogenetically distinct.

Amongst the additional genomes is the genome of TIGR4, which was the first pneumococcal strain to have its genome fully sequenced by The Institute for Genomic Research (TIGR, now J. Craig Venter Institute; accession number NC_003028.3) (16). Several differences were identified through alignment of the genome determined by TIGR, and the one presented here. The most significant are those in the repeat-rich genes *pavB* and *psrP* were deemed most significant. Within the TIGR genome, *pavB* is described to consist of four streptococcal surface repeats (SSUREs) (16). This contrasts to the five SSURE repeats we identify here. As for the number of repeats in *psrP*, a difference was observed in the second of the two serine-rich repeats (SRRs). The TIGR genome has been determined to have 539 (imperfect) repeats of the amino acid sequence SAS[A/E/V]SAS[T/I] in the SSR2 domain of PsrP, whereas we observe 225 (63). Besides the differences in these repeat-rich genes, no other changes in genetic content or structural differences were observed highlighting the high quality of the original TIGR4 genome sequence performed by shotgun Sanger sequencing.

## Discussion

In this work, we present an up-to-date iteration of the heavily used *S. pneumoniae* genome browser: PneumoBrowse 2. The continued development of novel methods to interrogate the *Streptococcus pneumoniae* genome is increasingly revealing the complex biology behind its regulation and organization. We have further refined the previous, already detailed, annotation of D39V by updating existing, adding new, and removing redundant features, but also by adding DNA methylation sites, sgRNA bindings sites, quantitative coupled transcriptome and proteome data, gene essentiality, chromosomal protein binding, and chromosome conformation data. Through the continued improvement of the annotation of its genomic content, the increasingly detailed annotation of D39V will continue to set the standard for the annotation of other genomes from both *S. pneumoniae* and other bacterial species.

Previous efforts to understand the regulation and contents of the *S. pneumoniae* D39V genome relied on RNA and DNA sequencing. However, these methods have not been able to fully determine the D39V proteome due to inherent challenges linked to transcript abundance, and computational algorithms. New work using ribosome profiling has led to the annotation of 122 previously undescribed sORFs, typically less than 50 amino acids in size (14). The small size does not mean that these sORFs cannot have a substantial impact on pneumococcal biology (64). For example, the competence-stimulating peptide of *S. pneumoniae* plays a crucial role in the regulation of competence and transformation, yet measures only 17 amino acids in length (65). As only eight sORFs have had their function (partly) presently elucidated, a vast potential for the discovery of novel biological mechanisms remains.

Besides updated D39V annotation, we extend the use of PneumoBrowse to a wider audience by providing the genomic sequence and annotations of 18 additional, phylogenetically diverse, *S. pneumoniae* strains, including that of TIGR4. The observed differences in two well-known repetitive regions (in *pavB* and *psrP*) between the TIGR4 genome determined by Tettelin *et al.* (16) and the one presented here, may either represent natural variation in closely related strains that has arisen through recombination events or mishaps during genome duplication or be the consequence of different genome sequencing techniques and assembly algorithms resulting in differences in these difficult-to-solve regions (66–68). Such variations have been noted before, as a different number of repeats in *pavB* of TIGR4 than the genome published by Tettelin *et al.*, have been reported before (69). These differences highlight the possible variability between different clones of the same strain, and the difficulties in solving the exact content of highly repetitive genomic loci despite long-read sequencing reads. These (and other) variations influence the exact genomic coordinates between these genomes.

Since the introduction of PneumoBrowse 1, other web-based resources that make the genome annotation of different *S. pneumoniae* strains available have also been developed. PneumoWiki (https://pneumowiki.med.uni-greifswald.de) provides information on coding features in the genomes of a wide variety of strains in a Wikipedia-style format, though it uses static webpages, and does not provide information on transcriptomics or nucleic acid binding by proteins. Other resources, not necessarily specific to *S. pneumoniae*, such as the KEGG database, the NCBI Nucleotide genome browser, or the BioCyc Genome Explorer, provide annotation information as well, but were developed with other specific goals in mind. The KEGG database for example, is aimed at linking genomic content to biological systems (such as biochemical pathways). The NCBI Nucleotide and BioCyc Genome Explorer genome browsers are a rich source of information but lack visualization of any additional information such as RNA quantification, or modified bases in the genome (39, 70, 71). PneumoBrowse however, offers a unique combination of visualization and interactivity. By combining the genome sequence, annotation, and a wide variety of other data types, PneumoBrowse stands out as a unique genome browser. As no database can provide the ever-growing range of information for every type of feature, we provide direct links to other databases, to complement the work available here, with the information presented there. We anticipate that the detailed characterization of D39V will lead to an increased understanding of this and other *S. pneumoniae* strains resulting in better preventive and treatment strategies for this clinically relevant opportunistic pathogen.

## Acknowledgments

We thank the Lausanne Genome Technologies Facility at the University of Lausanne, for PacBio sequencing, Colin Diesh and Doran Pauka for help with JBrowse 2. We thank Julien Dénéréaz, Clement Gallay, Xue Liu, and other members of the Veening lab for their valuable input and feedback. Strains 4954-98, DCC1476, GM17, M264-3, PJ755/1, SPN1439-106, R1501, SN75752, SN36255, SP264, TL7/1993, and TW31 were a kind gift of Mark van der Linden at the University Hospital RWTH Aachen. Strain EF3030 was a kind gift of Sven Hammerschmidt from the University of Greifswald. Genomic DNA of strain LILPNEUHC 19F was kindly provided by Jean-Claude Sirard from the Pasteur Institute in Lille. Genomic DNA of strain R1501 was kindly provided by Patrice Polard from the Center of Integrative Biology and the University Paul Sabatier in Toulouse. Strain BHN418 was a kind gift by Birgitta Henriques-Normark from the Karolinska Institute in Stockholm. Strain BM6001 and DP1322 were kindly provided by Francesco Ianneli from the University of Siena. We thank Michael J. Federle from the University of Illnois at Chicago, for the ribosome profiling data.

## Author Contributions

A.B.J. performed analyses, genomic DNA isolation and sequencing analyses, and wrote the manuscript with input from all other authors. P.S.G.: Performed genomic DNA isolation and sequencing analyses. A.M.B., V.d.B., and J.S. performed analyses. J.-W.V. oversaw the work.

## Funding

A.B.J. is supported through a Postdoctoral Fellowship grant (TMPFP3_210202) by the Swiss National Science Foundation (SNSF). P.S.G. was supported by a PhD fellowship of the Faculty of Biology and Medicine of the University of Lausanne. Work in the lab of J.-W.V. was supported by SNSF project grants (310030_192517 and 310030_200792), and the SNSF NCCR “AntiResist” program (51NF40_180541).

The funders had no role in study design, data collection and analysis, decision to publish, or preparation of the manuscript.

## Transparency declarations

The authors declare no competing interests.

## Notes

https://www.ncbi.nlm.nih.gov/bioproject/PRJNA1103744

### Competing Interest Statement

The authors have declared no competing interest.

